# MINN: A METABOLIC-INFORMED NEURAL NETWORK FOR INTEGRATING OMICS DATA INTO GENOME-SCALE METABOLIC MODELING

**DOI:** 10.1101/2025.05.14.653943

**Authors:** Gabriele Tazza, Francesco Moro, Dario Ruggeri, Bas Teusink, László Vidács

## Abstract

The understanding of cellular behavior relies on the integration of metabolism and its regulation. Multi-omics data provide a detailed snapshot of the molecular processes underpinning cellular functions and their regulation, describing the current state of the cell. While Machine Learning (ML) models can uncover complex patterns and relationships within these data, they require large datasets for training and often lack interpretability. On the other hand, mathematical models, such as Genome-Scale Metabolic Models (GEMs), offer a structured framework for analyzing the organization and dynamics of specific cellular mechanisms. At the same time, they don’t allow for seamless integration of omics information. Recently, a new framework to embed GEMs in a neural network has been introduced: these hybrid models combine the strengths of mechanistic and data-driven approaches, offering a promising platform for integrating different data sources with mechanistic knowledge. In this study, we present a Metabolic-Informed Neural Network (MINN) that utilizes multi-omics data to predict metabolic fluxes in *Escherichia coli*, under different growth rates and gene knockouts. We test its performances against pure ML and parsimonious Flux Balance Analysis (pFBA), demonstrating its efficacy in improving prediction performances. We also highlight how conflicts can emerge between the data-driven and the mechanistic objectives, and we propose different solutions to mitigate them. Finally, we illustrate a strategy to couple the MINN with pFBA, enhancing the interpretability of the solution.

## 1 INTRODUCTION

The phenotype of a cell is a complex interplay between its metabolic network, consisting of thousands of biochemical reactions, and the regulatory mechanisms controlling diverse cellular functions. Mechanistic models, such as GEMs, provide a structured framework to integrate and connect the available knowledge to find emergent properties in cellular systems. GEMs mathematically represent cellular metabolism, summarizing our information about the biochemical processes present in an organism. One common approach to simulate cellular behavior using GEMs is constraint-based modeling (CBM). Among these methods, Flux Balance Analysis (FBA) is particularly notable. FBA applies linear programming to optimize the distribution of metabolic fluxes, aiming to maximize specific objectives like biomass production while considering nutrient availability constraints. However, the predictive power of a mechanistic model like FBA is limited by the completeness of our understanding of cellular processes. Moreover, FBA typically have multiple feasible solutions. In such cases, the solution with the lowest sum of fluxes is usually selected, based on the assumption that cells try to minimize their enzyme production [Lewis et al., 2010]. However, this assumption is often an oversimplification, which does not account for the complex regulatory mechanisms within cells.

On the other hand, data-driven machine learning (ML) models can effectively extract patterns in high-dimensional datasets, such as multi-omics data, without prior knowledge of underlying molecular mechanisms. These models often demonstrate strong predictive capabilities, but are limited by the scarcity of biological datasets, frequently constrained by experimental costs and time. In recent years, ML models have also been explored for predicting metabolic fluxes using multi-omics data. Although FBA remains the preferred mechanistic framework for this task, integrating omics data into CBM frameworks remains a significant challenge (Machado and Herrgård [2014]). Interestingly, recent work [Gonçalves et al., 2023] demonstrated that purely ML-based approaches trained on omics data can outperform FBA-based methods in metabolic flux prediction.

Given the complementary strengths and limitations of ML and GEMs, there has been increasing interest in trying to combine these two approaches (Zampieri et al. [2019]). Hybrid models that merge mechanistic knowledge with the predictive capabilities of ML offer a promising direction. Recently, Faure et al. [2023] developed a framework that combines CBM with ML in a neural network architecture called an Artificial Metabolic Network (AMN). Their approach leveraged GEM structures and FBA constraints within neural networks to predict growth rates from media compositions.

In Faure et al. [2023], the authors suggested three different possible configurations for incorporating the GEM structure and the FBA constraints in a neural network (NN). In this work, we selected one of these configurations, inspired by Physics-Informed Neural Networks (PINNs), and we re-implemented and expanded it to integrate multi-omics data as inputs. We will refer to this new architecture as Metabolic-Informed Neural Network (MINN) with multi-omics integration (Figure 1). We applied this hybrid model to the dataset analyzed by Gonçalves et al. [2023], which examines how the metabolism of *Escherichia coli* adapts to varying growth rates and single-gene knockouts (Ishii et al. [2007]). As discussed in Tazza et al. [2024], the combination of fluxes measured experimentally lies outside the solution space of FBA, causing a conflict between the optimization of the data-driven and the mechanistic objectives. To address this, we provide different mitigation strategies.

**Figure 1:**
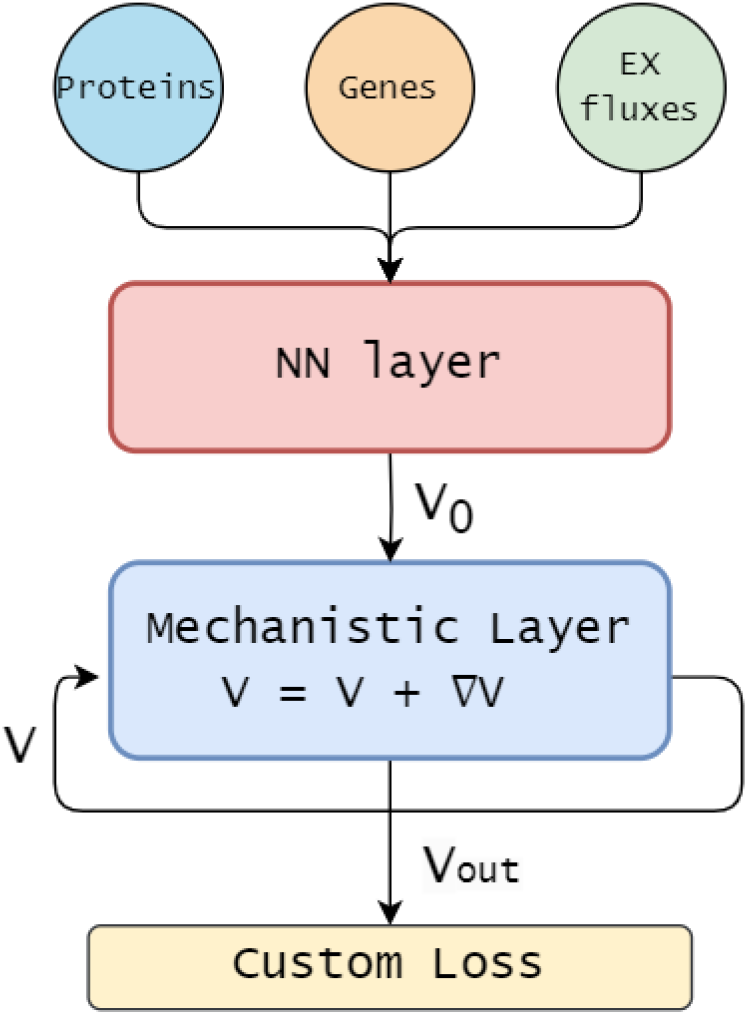
Schematic representation of the MINN architecture. Protein, gene, and exchange flux data are used as input to a feed-forward neural network, which produces an initial estimate for the flux distribution *V*_0_. This estimate is refined in a mechanistic layer via a gradient descent step to better align with flux balance constraints, resulting in the final flux distribution *V*_*out*_. A custom loss function combining data error and FBA constraint violation is used to train the network using backpropagation.

To summarize, in this work:

- We describe the implementation of a MINN with multi-omics integration, using an early concatenation approach.
- We benchmark its predictive performances on the ISHII dataset, compared to pure ML methods Gonçalves et al. [2023].
- We recalculated the data to be in the FBA solution space and compared the predictive performances with those on the original measurements.
- We explore different hybrid optimization strategies to address the conflict between the objectives, while using the original data.
- Finally, we adapt the MINN to a “reservoir” configuration Faure et al. [2023], which uses the MINN predictions to directly constrain pFBA, and compare its predictions with those of pFBA alone.

With these analyses, we provide a detailed overview of methods and strategies to adapt and use hybrid ML-FBA methods for multi-omics integration. Our findings highlight the potential of hybrid models to enhance the predictive accuracy and robustness of metabolic flux predictions. This paves the way for more precise and comprehensive metabolic network analyses, particularly for phenotypes where metabolism is significantly influenced by other layers of cellular organization, which are challenging to incorporate into FBA. Furthermore, with this work we aim to provide a guide to the use of the MINN framework, helping researchers choose the most suitable configuration based on the specific objective of their study.

## 2 RELATED WORKS

### 2.1 Genome-Scale Metabolic Models and Flux Balance Analysis

A genome-scale metabolic model (GEM) is a comprehensive reconstruction of an organism’s metabolic network, representing the full metabolic capacity encoded by its genome. It serves as a structured knowledge base, integrating information on genes, proteins, enzymes, and metabolic pathways [Monk et al., 2014, O’Brien et al., 2015]. GEMs have been primarily reconstructed for microorganisms, but models also exist for multicellular organisms, including humans. A typical microbial GEM contains hundreds or even thousands of reactions and metabolites, increasing in complexity for multicompartment systems like yeast [Somerville et al., 2022]. To analyze such large models, a commonly used method is Flux Balance Analysis (FBA), which relies on the assumption of steady state or balanced growth [Orth et al., 2010a, Bruggeman et al., 2020]. Under these conditions, the concentrations of metabolites remain constant over time, and the rates of production and consumption are balanced across all reactions. GEMs can therefore be formulated only in terms of reactions rates and treated as linear programming problems. FBA uses this framework to predict the metabolic behavior of the organism by optimizing, given the stoichiometric constraints, a specific objective function: commonly biomass production or, in biotechnological contexts, the yield of a desired product. However, stoichiometric constraints alone are often insufficient to determine a realistic flux distributions. To improve predictive accuracy, FBA requires context-specific inputs—such as growth medium composition—typically incorporated by constraining exchange fluxes. These constraints may be derived from experimental measurements, assumptions about nutrient uptake kinetics, or a combination of both [Palsson, 2015].

### 2.2 Omics Data Integration in Genome-Scale Metabolic Models and Flux Balance Analysis

GEMs are reconstructed primarily from genomic information, encoding the metabolic network of an organism. A GEM can be tailored to represent different cell strains by including or excluding reactions based on the presence or absence of genes encoding the relevant metabolic enzymes [Thiele and Palsson, 2010]. Beyond genomics, a broad range of omics data (e.g., transcriptomics, proteomics, metabolomics, and fluxomics) can significantly enhance the accuracy and predictive capacity of GEMs [Machado and Herrgård, 2014, Yizhak et al., 2010]. The challenge lies in translating these data into metabolic fluxes. For enzyme-catalyzed reactions, the reaction rate is typically described by:

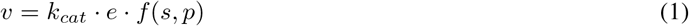

where:

- *v* is the reaction’s flux;
- *k*_*cat*_ is the catalytic constant, or turnover number, which represents the number of substrate molecules converted to product per enzyme molecule per unit time when the enzyme is fully saturated with substrate, i.e. the enzyme efficiency;
- *e* is the enzyme concentration;
- *f* (*s, p*) is a (often nonlinear) function of the concentrations of substrates *s* and products *p*, and the corresponding affinity parameters.

It is worth also noting that, while genomic, transcriptomic, and proteomic data provide rich layers of information, only features that can be explicitly linked to metabolic reactions can be directly integrated into GEMs [Liebermeister et al., 2014]. As a result, much of the broader cellular context captured by these datasets, including regulatory, structural, or signaling components, is typically excluded from the model.

#### Transcriptomics data

while often weakly correlated with actual protein levels or enzymatic activity [Ponomarenko et al., 2023], remain useful to identify active metabolic genes and infer condition-specific pathway activation. When comparing multiple conditions, transcriptome profiles may suggest shifts in metabolic strategy. Several frameworks have been developed to incorporate transcriptomics and gene co-expression data into GEMs [Machado and Herrgård, 2014, Paklao et al., 2023, Zampieri et al., 2023].

#### Proteomics data

provide a closer proxy for metabolic capability. Presence or absence of specific enzymes can directly constrain which reactions are allowed under given conditions. Assuming enzyme saturation (*f* (*s, p*) = 1), enzyme levels scaled by *k*_*cat*_ values provide upper bounds for fluxes (*v*_*max*_). These catalytic parameters can be obtained from databases such as BRENDA [Chang et al., 2021] and SABIO-RK [Wittig et al., 2018], or estimated using statistical approaches or enzyme-kinetic models [Singh et al., 2024, Diamataris et al., 2023]. Although these resources are growing rapidly, retrieving the relevant kinetic parameters remains a semi-automated process that often requires manual curation to ensure accuracy and model compatibility. Building on this concepts, more complex model formulations have been developed, such as enzyme-constrained GEMs (ecGEMs), which treat metabolism as a problem of protein budgeting under limited cellular capacity [Chen et al., 2024, Salvy and Hatzimanikatis, 2020]. Proteome-constrained models (pcGEMs) further account for the resource cost of enzyme production [Grigaitis et al., 2021, Elsemman et al., 2022]. While building these models often requires custom pipelines, tools such as GECKO [Chen et al., 2024] and sEnz [van den Bogaard et al., 2024] are making these tasks increasingly standardized and accessible.

#### Metabolomics data

offer insights into both the structure and dynamics of metabolism. Although metabolite concentrations cannot be directly used in GEMs due to the steady-state assumption and lack of proportionality between concentrations and fluxes, their presence or absence can indicate pathway activity. Time-series measurements of extracellular metabolite levels can be converted into flux constraints for exchange reactions, allowing tuning the GEMs to match observed uptake or secretion patterns Henriques et al. [2021]. If thermodynamics is included, the metabolic data can also be used to constrain or validate fluxes [Niebel et al., 2019].

#### Fluxomics data

isotope-labeling experiments (e.g., growth in ^13^C-glucose medium) allow direct estimation of intracellular flux distributions, via Metabolic Flux Analysis (MFA) [Antoniewicz, 2021]. These experimentally derived fluxes can be used to constrain specific reactions or to find the flux profile that best fits the measured data.

#### Multi-omics data

The simultaneous integration of diverse omics layers yields a more holistic view of the cellular state. This systems-level approach is especially valuable for understanding dynamic or context-dependent responses, as it captures the interactions between different biological processes [Lu et al., 2018]. Tools like the IOMA (Integrative Omics-Metabolic Analysis) framework facilitate the incorporation of diverse omics layers into GEMs to enhance predictive fidelity [Yizhak et al., 2010].

In summary, omics data integration represents a powerful way to contextualize and refine GEMs, improving their ability to simulate real-world biological behavior. However, current methods remain limited by a lack of standardization and automation. Moreover, mechanistic integration is restricted to metabolic features explicitly represented in GEMs, meaning that valuable context from a broader view of cellular processes is still usually excluded.

### 2.3 Integrating FBA and Machine Learning for Enhanced Metabolic Predictions

In the previous sections, we discussed how FBA is a powerful approach to exploit the information stored in GEMs to predict the metabolic behavior of cells. However, FBA has at least four main limitations. First, its predictive power heavily depends on the amount of experimental measurements of exchange fluxes. Second, incorporating multi-omics data is challenging, because all measurements must be converted into fluxes, a process that usually requires iterative steps of time-consuming manual curation. Third, FBA and GEMs focus solely on metabolism and typically do not link it to the general status of the cell. Finally, FBA predict flux distributions that tend to maximize the yield on the limiting substrates [van Pelt-KleinJan et al., 2021], often missing to capture “high rate-low yield” solutions [Elsemman et al., 2022].

In recent years, with the increasing availability of high-throughput technologies and data, ML has gained popularity as a valid alternative to mechanistic-based approaches [Wytock and Motter, 2018, Gonçalves et al., 2023, Al et al., 2024]. The success of ML lies in its ability to find patterns in the data without making any mechanistic assumptions. However, a main drawback is that ML requires a high volume of data to train models successfully, and in many biology-related domains, datasets of suitable size are rare. In particular, experiments in microbial physiology tend to be one, if not two, orders of magnitude smaller than what ML requires. Moreover, ML behaves mostly as a black-box model, making it difficult to extract mechanistic understanding from its results. On the other hand, this black-box nature makes ML more amenable than mechanistic models for integrating diverse data sources, even those for which there is no clear understanding of their connections.

Therefore, it seems natural to integrate these two approaches to overcome each other’s limitations and exploit their strengths. In recent years, as reviewed in Sahu et al. [2021] and Zampieri et al. [2019], there have been many attempts to integrate these methods. Sahu et al. [2021] categorize these works into two groups: ML as input of FBA Dai et al. [2018], Kim et al. [2016], Morrissey et al. [2025] and FBA as input of ML Magazzù et al. [2021], Culley et al. [2020]. This division highlights that these methods do not truly integrate ML and FBA but rather concatenate them, using them in two distinct steps. To the best of our knowledge, only two works presented hybrid models that genuinely integrate FBA and ML: Faure et al. [2023] and Hasibi et al. [2024]. The first introduces Artificial Metabolic Neural Networks (AMNs), which are Neural Networks that use FBA constraints to refine their solution and to regularize the network. This is achieved through a Mechanistic Layer, representing the structure of the mechanistic model inside the NN, and a custom loss function, similar to those of other Knowledge Informed Neural Networks (e.g. Physics-Informed Neural Network [Cuomo et al., 2022]). When using FBA alone for growth rate prediction, nutrient uptake fluxes often need manual adjustment to match experimental growth rates. This process can involve labor-intensive experiments or unsystematic “trial-and-error” adjustments, which may introduce arbitrary assumptions to align the model with observed data. The hybrid AMN framework proposed by Faure et al. [2023] addresses these challenges by embedding mechanistic information into neural networks, providing a more systematic approach. The latter presents FlowGAT, which integrates the structure of the GEM and the solution of FBA in a Graph Attention Network (GAT) to predict the genes essentiality.

The MINN models presented here follow the blueprint of AMNs and, in line with approach presented in Faure et al. [2023], represent a true hybrid model, integrating FBA constraints and multi-omics data to improve predictions of fluxes. A first implementation of the model was presented in Tazza et al. [2024]. Here we further develop this architecture and test different solutions to mitigate conflicts that emerged between the data-driven and the mechanistic objectives.

## 3 METHODS

### 3.1 Dataset

The dataset analyzed in this work was originally published by Ishii et al. [2007] and consists of 29 chemostat experiments, in which *E. coli* was grown in glucose minimal medium. Wild-type strain K-12 was grown at 5 different dilution rates (D = 0.1, 0.2, 0.4, 0.5, and 0.7 h^−1^), while 24 different single-knockout mutant strains were cultivated at fixed dilution rate (D = 0.2 h^−1^). The same dataset was already used by Gonçalves et al. [2023] to test traditional ML for the prediction of metabolic fluxes from multi-omics data.

The dataset consist of transcriptomic, proteomic, and fluxomic measurements. For each sample, microarrays were used to assess the expression profiles of 79 genes and LC-MS/MS quantitative proteomics to measure the abundances of 60 proteins. ^13^C-labeled metabolomics experiments were analyzed with MFA to estimate 47 metabolic fluxes: 37 reactions of the central carbon metabolism, 9 exchange fluxes (production or consumption of external metabolites) and biomass growth.

The metabolic model used by Ishii et al. [2007] to perform MFA is a core model that mainly represents the central carbon metabolism of *E. coli* and how it connects to the measured external metabolites. This model is much smaller and less complete than the GEM [Feist et al., 2007] integrated in the MINN. For the GEM to grow, many more different biomass components must be synthesized, diverting some metabolic precursors outside the pathways represented in the MFA model. For this reason, the fluxomics data from Ishii et al. [2007] lie outside the solution space [Tazza et al., 2024]. In most of our analyses we used the original fluxomics data, to highlight the ability of the MINN to reconcile MFA fluxomics data with the structure of the full-size GEMs. However, to investigate the impact of this discrepancy, we repeated some of the analyses with a second set of fluxes, now residing in the FBA solution space. This second set of fluxomics data is composed of the fluxes with the minimum Euclidean distance from the original ones, following an approach detailed in the supplementary material of Machado and Herrgård [2014] and we refer to it as *FBA fit* data.

### 3.2 GEM preparation

In this section we describe all the genome-scale metabolic reconstructions utilized to build the MINNs. The most recent GEM available for E. coli K-12 is iML1515 [Monk et al., 2017], but we opted for iAF1260 [Feist et al., 2007]. The two differ mainly for the more comprehensive coverage of accessory pathways of iML1515, which are relevant in complex environments like the human gut, but not for growth on minimal medium. On the other hand, the size of the GEM can heavily affect the complexity of the MINN: using a smaller GEM would improve the efficiency of our hybrid model by reducing the computational resources required for training. Possibly, it would also reduce the noise in the model, enhancing the prediction accuracy. iAF1260 is reasonably smaller than iML1515 (2382 reactions vs. 2712) and is also the same model used by Gonçalves et al. [2023] in their analyses.

We further reduced the size of the model by excluding all the reactions which cannot carry flux during growth in glucose minimal medium. This was achieved performing Flux Variability Analysis (FVA) and retaining only the reactions with a non-zero span. We refer to this model as *FVA-reduced* GEM. We also tested a second strategy, inspired from Faure et al. [2023], to further reduce the model. We generated a dataset of 2000 FBA solutions by randomly selecting single-gene knockouts and varying the maximum glucose uptake rate within the experimentally observed range. Reactions that consistently carried zero flux across all the simulations were removed from the model. We refer to this model as *FBA-reduced* GEM. To investigate the impact of an extreme decrease in the genome-scale reconstruction size, we also built a MINN using the e_coli_core model [Orth et al., 2010b], a manually reduced GEM focused on central carbon metabolism, which is the smallest model available in the BiGG database.

Finally, to further test the role of the GEM and the underlining metabolic network, we also tested our baseline configuration including the GEM of a different organism. We used the iNF517 model Flahaut et al. [2013] for *Lactococcus lactis subsp. cremoris MG1363*. This microorganism is a lactic acid bacterium, with an incomplete TCA cycle, which makes it an interesting comparison for *E*.*coli*, both in terms of structure of the network in the central carbon metabolism and of general metabolic behavior. The model was reduced using the FVA-guided reduction approach. The results of this comparison are available in the Supplementary Materials.

In Table 1 we summarize the dimensions of each GEM. The GEMs were downloaded from the BiGG database (King et al. [2016]) and handled/modified using CBMPy 0.8.4 (Olivier et al. [2023]). In each model, reversible reactions were split into a forward and a reverse reaction using the built-in CBMPy function `cbmpy.CBTools.splitReversibleReactions`.

**Table 1:**
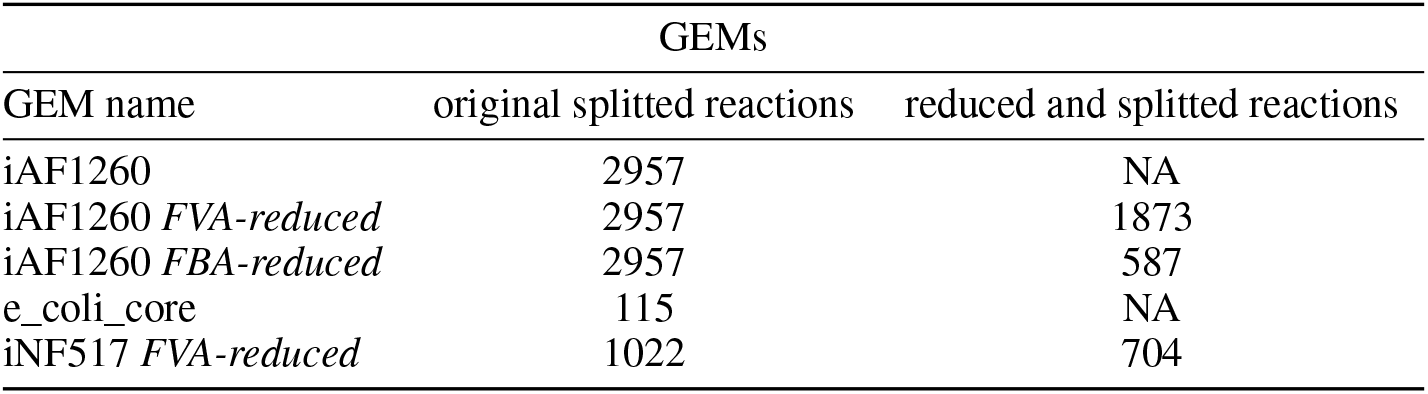
Dimensions of all the GEM used in this analysis.

| GEM name | GEMs |  |
| --- | --- | --- |
|  | original splitted reactions | reduced and splitted reactions |
| iAF1260 | 2957 | NA |
| iAF1260 *FVA-reduced* | 2957 | 1873 |
| iAF1260 *FBA-reduced* | 2957 | 587 |
| e_coli_core | 115 | NA |
| iNF517 *FVA-reduced* | 1022 | 704 |

### 3.3 MINN architecture

This study introduces a MINN architecture designed to predict multiple fluxes using multi-omics data, which provide key insights for metabolic predictions but are challenging to integrate with FBA (Machado and Herrgård [2014]). Figure 1 illustrates the structure of our MINN architecture, built to predict fluxes measured in the ISHII dataset using proteomics, transcriptomics, and the measurements of two exchange fluxes, namely (*R_EX_glc D_e, R_EX_o2_e*). The data are integrated using an early concatenation strategy Adossa et al. [2021], where the three omics datasets are combined into a single matrix that is fed into the MINN.

The first part, red in Figure 1 is a pure feed-forward NN where the input dimension is equal to the number of data features *d*_*in*_ and the output dimension equal to the number of the total reactions in the GEM, *d*_*out*_. The output *V*_0_ represents the NN’s initial guess for the flux distribution. The second part, the mechanistic layer, blue in Figure 1, consists of a gradient descent optimization loop. This loop refines the output of the first neural network step by adjusting the final predicted flux distribution *V* to better comply with the FBA constraints, minimizing the FBA loss function *L*_*F BA*_. The neural network weights are trained using a standard back-propagation algorithm and a custom loss function that considers both the data error and the FBA constraints *L*_MINN_.

To formalize this, let the input data be 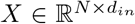 where *N* is the mini-batch dimension in the back-propagation algorithm and *d*_*in*_ is the number of features of *X*. The first NN part can be expressed as:

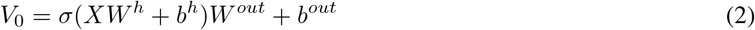

with *σ* the ReLU activation function, 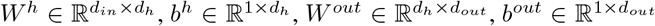, the weights matrices and the biases of the input and hidden layer respectively, where *d*_*h*_ is the hidden layer dimension and *d*_*out*_ is the output dimension, which coincides with the dimension of the flux distribution. Then, *V*_0_ is refined in the mechanistic layer through a gradient descent optimization. For simplicity, the notation refers to the simple case when the loop has one iteration:

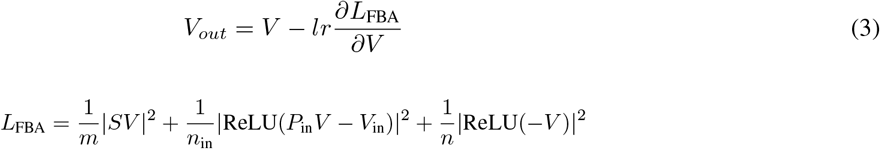

The first element represents the steady-state constraint of FBA, with *S* as the stoichiometric matrix of the GEM. The term *m* denotes the number of metabolites and serves as the normalization term. The second element represents the upper-bound constraint on the vector of fluxes *V*_*in*_. Here, *P*_*in*_ represents the projection matrix that projects the flux distribution vector *V* into the dimension of *V*_*in*_, while the normalization term *n*_*in*_ stands for the number of bounded fluxes. Lastly, the last element symbolizes the lower bound constraint, which required since the GEM is built to ensure that all the fluxes are positive.

The custom loss used to train the weights of the MINN is:

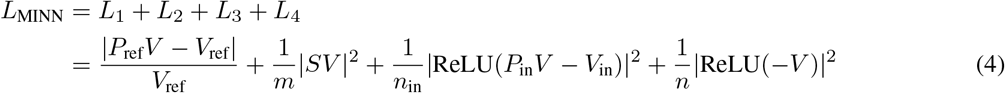

where *V*_*ref*_ is the vector with the measured fluxes and *P*_*ref*_ a projection matrix that projects *V* to the dimension of *V*_*ref*_ ; while the other elements represent the FBA constraints and are the same of *L*_*F BA*_.

In Faure et al. [2023], the authors used Mean Squared Error (MSE) as *L*_1_ because they wanted to predict a single flux, specifically the growth rate. In our work, we use the Normalized Error (NE) [Gonçalves et al., 2023] to have a scale-invariant *L*_1_ when predicting multiple fluxes in order to avoid favoring reactions with higher flux values.

In addition, during our analysis a conflict between the data-driven and mechanistic losses emerged. In order to mitigate this issue, we multiply *L*1 with a constant *c*, which allow us to adjust the balance between the two losses:

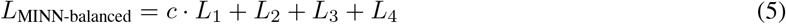

The *c* constant becomes a hyperparameter of the model, tuned using k-fold cross-validation and the optimized value determines the best balance between *L*_1_ and (*L*_2_ + *L*_3_ + *L*_4_).

#### 3.3.1 MINN-reservoir

Similarly to Faure et al. [2023], we tested an additional configuration of the MINN, named MINN-reservoir. The training of the MINN-reservoir requires two steps, as shown in Figure 2. In the first step (2a), a MINN (with no omics data in input) is trained only to reproduce FBA using a dataset of FBA solutions. This dataset contains the results of 2000 FBA simulations, in which the reactions belonging to *V*_*in*_ *(R_EX_glc D_e, R_EX_o2_e, R_EX_co2_e, R_EX_etoh_e and R_EX_ac_e)* were assigned random values, within the ranges of variability observed in the ISHII dataset. This procedure creates a model that acts as a pure approximator of an FBA solver (Pretrained block in Figure 2), capable of predicting the optimal flux distributions from measurements of external metabolites fluxes, similar to an FBA solver. In the second step (Figure 2b), this Pretrained block is embedded into a new MINN architecture, where it replaces the Mechanistic Layer. The resulting architecture consists of a neural network layer that predicts *V*_*in*_ from multi-omics data and medium exchange fluxes (*R_EX_glc D_e, R_EX_o2_e*), followed by the *Pretrained block*, which computes the flux distribution *V*_*out*_ from the predicted *V*_*in*_. For the test sample in each split, the predicted *V*_*in*_ are extracted and used as additional constraints for pFBA: increasing the input information in a data-driven way while preserving the mechanistic structure of the model. Unlike the default configuration of the MINN, this approach produces as final output not only the full flux distribution, but a complete solution from a Linear Programming solver, which can be analyzed with all the tools developed for this purpose.

**Figure 2:**
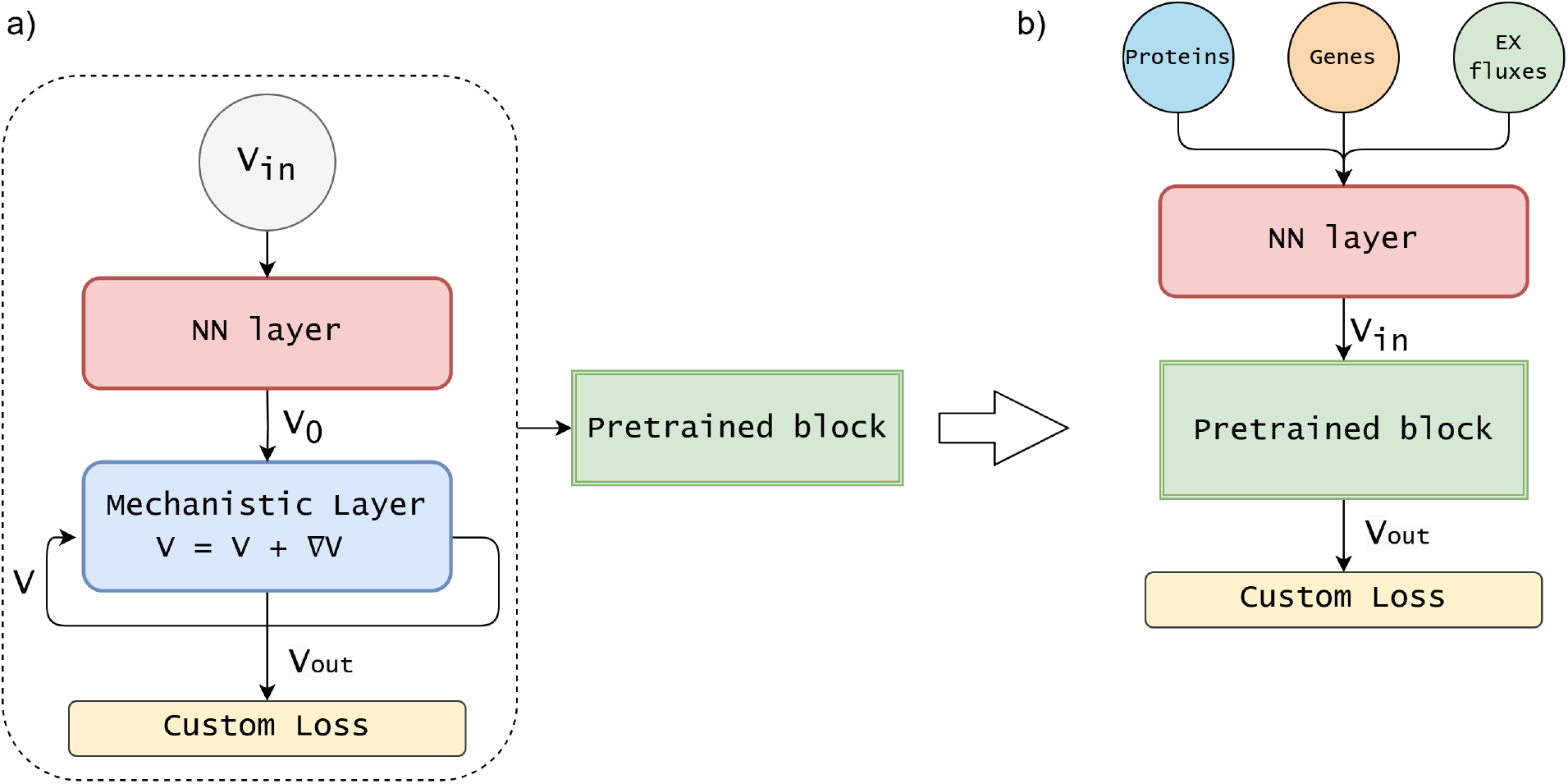
Two-step training strategy of the MINN-reservoir architecture: a) In the first step, a MINN (with no omics data in input) is trained to approximate an FBA solver, using a dataset of simulated FBA solutions. The network learns to predict the flux distribution *V*_*out*_ from randomly sampled external fluxes *V*_*in*_ *(R_EX_glc D_e, R_EX_o2_e, R_EX_co2_e, R_EX_etoh_e and R_EX_ac_e)*. Once trained, its weights are frozen, and the resulting model is reused as a fixed *Pretrained block*. b) In the second step, this *Pretrained block* is embedded within a new architecture that takes omics data and medium exchange fluxes (*R_EX_glc D_e, R_EX_o2_e*) as input. A neural network predicts *V*_*in*_, which is then passed to the *Pretrained block* to compute *V*_*out*_.

#### 3.3.2 Hybrid Optimization Strategies for Data-Driven and Mechanistic Integration

The equation 4 highlights how the loss of the MINN is composed of two components: *L*_1_, which drives the optimization on the data, and *L*_*F BA*_, which minimize the divergence from the mechanistic model. In Equation 5, we already introduced the coefficient *c*, which allows to tweak the balance between the two components, either manually or through hyperparameter optimization. In this section, we introduce three other methods to tune this balance while minimizing the trade-off between the different components.

In developing hybrid models that integrate mechanistic constraints with data-driven approaches, we introduce several methods to balance the objectives of maintaining adherence to the FBA constraints without substantially compromising predictive performance. These methods address the challenge posed by different scales of mechanistic and data-driven losses, ensuring that neither dominates the optimization process and that the model generalizes well to unseen data.

##### Bound on Mechanistic Loss

The first method introduced is a bound on the mechanistic loss. This method ensures that the model’s predictions do not deviate too far from the mechanistic solutions. A fixed threshold is set, and when the mechanistic loss exceeds this threshold, a multiplicative factor is applied to penalize further deviations. This approach softly constrains the model within a feasible solution space derived from mechanistic constraints, preventing the model from paying an excessive cost in terms of mechanistic loss to improve the data driven. To provide a clear view of the described bound on the mechanistic loss, a graphical representation is available in the Supplementary Material file.

##### Loss Balancing

A loss balancing mechanism was employed to handle the different scales of mechanistic and data-driven losses. This method normalizes each loss by dividing it by the exponential average of its previous values, as detailed in Hu et al. [2018]. Through this approach, both losses are considered equally during gradient updates, preventing one from outweighing the other during the training process. The loss balancing ensures that the mechanistic loss, which would typically be underrepresented due to its smaller scale, contributes adequately to model optimization alongside the data-driven loss.

##### Loss Weight Scheduler

Lastly, a dynamic loss weight scheduler was implemented to gradually shift the model’s focus between the mechanistic and data-driven tasks throughout training. For the first phase, the scheduler prioritizes the mechanistic loss, ensuring it starts from a solution closer to the mechanistic model’s feasible space. As training progresses, there is a transition phase where the scheduler gradually increases the importance of the data-driven loss until it reaches the final phase, where the data-driven loss has a higher weight, guiding the model toward better predictive performance for the data-driven task. For the sake of clarity, a graphical representation of the weight scheduler is available in the Supplementary Material file.

These methods provide a comprehensive framework for balancing the trade-offs between data-driven accuracy and mechanistic integrity.

### 3.4 Computational setup

We adopted the same evaluation pipeline for all our MINN configurations to evaluate the MINN performance and have a fair comparison with the results obtained by [Gonçalves et al., 2023]. It consists of a dual-loop cross-validation process. The outer loop is a leave-one-out, and in each train loop, there is an inner loop of a k-fold with *k* = 5 to tune the hyper-parameters. The tuning concerns the dimension of the first hidden layer, the learning rate of the NN, the intensity of dropout and *L*2 regularization, and the *c* constant for the *L*_1_ loss in the case of the MINN-c-balanced. For a consistent comparison with the work of Gonçalves et al. [2023], we employed identical metrics to evaluate all our experiments: the regression coefficient *R*^2^, the mean absolute error (MAE), the root mean squared error (RMSE) and the normalized error (NE). As described later, in some of our results, we also report the *L*_2_ as a metric to measure the quality of the predicted flux distribution.

## 4 RESULTS AND DISCUSSION

This work aims to compare the predictive performance of the MINN w.r.t pure ML approaches and mechanistic models such as pFBA. For clarity, we divide all the results and discussions into three groups. The first one (Table 2) contains the performance comparison results between our approach and the pure ML ones presented in Gonçalves et al. [2023]. Here, we evaluate the predictive performance of different methods on the 45 reference fluxes measured in the ISHII dataset. The second one (Table 3) includes the results of the comparison between different approaches employed to mitigate the issue of conflicting losses. Here, we compare the performance of the different methods on the measured fluxes and the quality of the predicted flux distribution, which we measure using, as a proxy, *L*_2_. The third group (Table 4), instead, compares the results obtained using the MINN-reservoir configuration with those of pFBA.

**Table 2:**
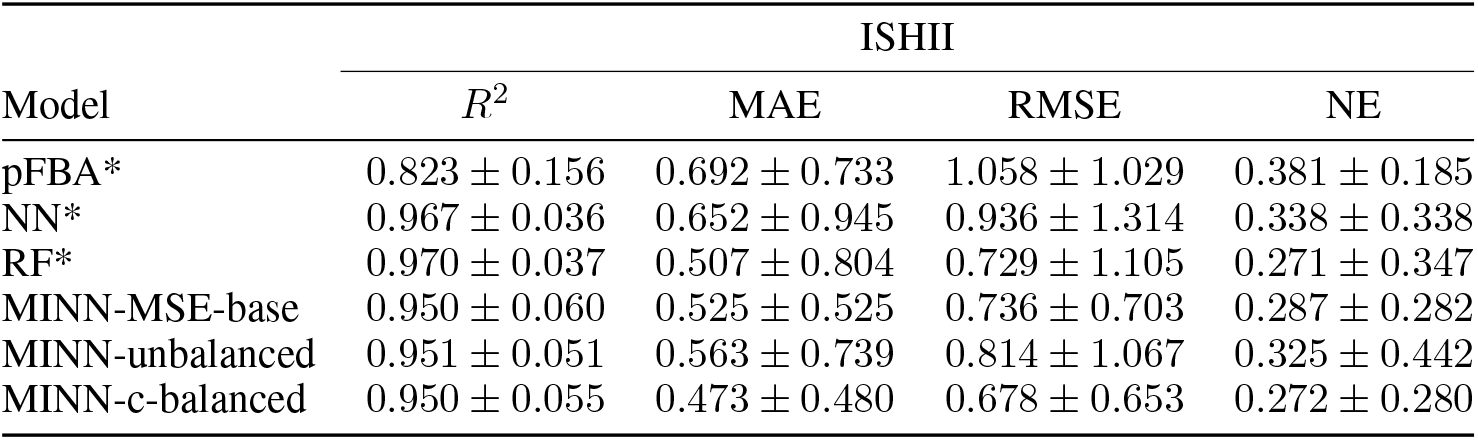
Comparison of predictive performance between our proposed MINN-based approaches and purely mechanistic and machine learning methods from Gonçalves et al. [2023]. Metrics average and standard deviation over 29 leave-one-out splits. *results from Gonçalves et al. [2023]

**Table 3:**
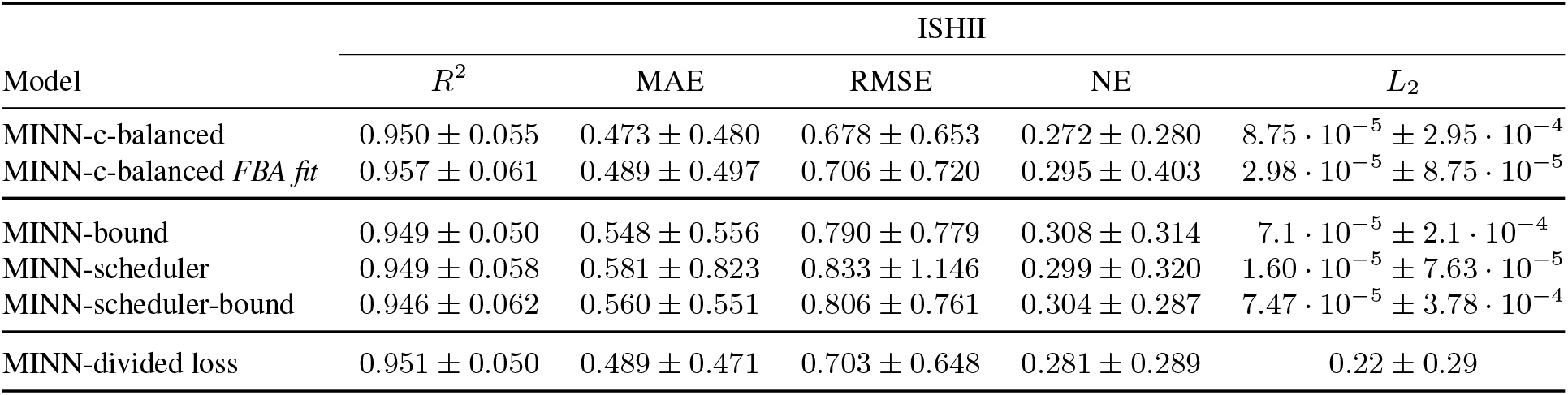
Performance comparison of different methods addressing the issue of conflicting losses. Metrics average and standard deviation over 29 leave-one-out splits.

**Table 4:** Performance comparison between the standard pFBA and MINN-reservour + pFBA. The results evaluate the effectiveness of the MINN-reservoir approach to enrich the input for pFBA in comparison to standard pFBA. Metrics average and standard deviation over 29 leave-one-out splits.

| Model | ISHII |  |  |  |
| --- | --- | --- | --- | --- |
| | $R^2$ | MAE | RMSE | NE |
| pFBA | $0.892 \pm 0.127$ | $0.496 \pm 0.353$ | $0.836 \pm 0.625$ | $0.306 \pm 0.367$ |
| MINN-reservoir + pFBA | $0.910 \pm 0.091$ | $0.445 \pm 0.261$ | $0.740 \pm 0.421$ | $0.253 \pm 0.159$ |

Lastly, we tested the impact of using different GEMs in the mechanistic layer of the MINN. In particular, when we included the GEM of *Lactococcus lactis* subsp. *cremoris*, a bacterium with an incomplete TCA cycle compared to *E. coli*, we observed a decrease in the quality of the predicted flux distribution. Although the difference was moderate, probably due to the conservation of central carbon metabolism, these results suggest that the biological relevance of the GEM has an important role in the regularizing effect of the mechanistic layer. A more detailed analysis is available in the Supplementary Materials.

### 4.1 MINN to predict measured fluxes

In the first group, as a baseline, we used the MINN architecture with an MSE as *L*_1_ (MINN-MSE-base), as in Faure et al. [2023]. As shown in Table 2, the results are already comparable with a Random Forest (RF), the best machine learning method in Gonçalves et al. [2023], and better than the NN approach. To avoid potential bias from the large discrepancies (up to two orders in magnitude) between the values of the fluxes we are predicting, we replaced the MSE in *L*_1_ with a NE. As detailed in the Section 3, we multiply *L*_1_ with a constant *c*, optimized during the cross-validation. This method, incorporating the *c* parameter, is referred to as MINN-c-balanced, while the one without this adjustment as MINN-unbalanced. Although the change from MSE to NE ensures a scale-invariant *L*_1_, it also reduces its magnitude, amplifying the conflict between losses as the mechanistic constraints gain more influence. For this reason, the MINN-unbalanced obtained worse results w.r.t. the MINN-MSE-base, but the MINN-c-balanced shows the best results, achieving comparable or better performance than the RF in three out of four metric averages.

Moreover, the MINN-c-balanced shows a relevant reduction in standard deviation. This suggests that the inclusion of biological constraints stabilizes the learning process, reduces overfitting, and leads to more robust and consistent predictions across the 29 leave-one-out splits. The results confirm that MINN is a state-of-the-art approach for flux prediction, outperforming pure ML methods, like RF, and purely mechanistic approaches, like pFBA. Moreover, MINN not only learn flux values from data but also derive the entire flux distributions, as done by FBA simulations.

The challenge of predicting multiple fluxes, with significant variability in their values, was effectively addressed by substituting the MSE with an NE for the *L*_1_ and adding the *c* constant that handled the imbalances emerged. The MINN’s flexibility also makes it a powerful tool for integrating multi-omics and potentially other kinds of data with GEMs, enabling its application in diverse systems biology contexts.

### 4.2 MINN to predict a qualitative flux distribution

Regarding the second group of results, we aim to compare different methods to mitigate the issue of the conflicting losses already introduced in the Section 3. We address this problem for two different points of view. First, we act on a mechanistic aspects of the MINN: the reference data. In contrast, the second perspective addresses the optimization process of the MINN, where we apply various hybrid optimization strategies discussed in Section 3.3.2 As a baseline, we employ our best method in terms of prediction performance, hence MINN-c-balanced.

In the first case, we compare it with a configuration of the same MINN-c-balanced which uses different fluxes data, recalculated to be in the solution space of FBA. We described this process in 3.1 section. As shown in the first section of Table 3, this approach, named MINN-c-balanced *FBA fit*, does not reduce the quality of the fluxes prediction, represented by the four metrics, but it improves the*L*_2_ by reducing its value by a third.

The results show that MINN maintains strong predictive performance even when the flux data are not in the FBA solution space. This suggests that its data-driven component can compensate for deviations from mechanistic constraints. However, using flux data that aligns with the FBA solution space improves the quality of the predicted flux distribution. This correction makes the optimization process easier by alleviating the issue of conflicting losses.

Regarding the hybrid optimization strategies, we compare the MINN-c-balanced with three other methods previously introduced in Section 3.3.2: *MINN-bound*, which penalizes violations of the mechanistic loss that exceed a threshold; *MINN-scheduler*, which gradually shifts the focus from mechanistic to data-driven loss during training; and *MINN-scheduler-bound*, which combines both approaches. From the second section of the table, we observe that the *MINN-c-balanced* model achieves the lowest RMSE, indicating the best performance on the data-driven task. However, this comes at the cost of a higher *L*_2_, suggesting a trade-off where improved performance is achieved at the expense of mechanistic fidelity. On the other hand, the models incorporating a mechanistic bound, such as *MINN-bound* and *MINN-scheduler-bound*, show a small increase in RMSE and a modest reduction in *L*_2_, suggesting a limited effect on improving the trade-off between mechanistic accuracy and data-driven performance. Interestingly, the *MINN-scheduler* model finds a better compromise between these objectives: at the cost of a moderate RMSE worsening, it achieves the lowest *L*_2_ by a consistent margin. This demonstrates the strength of dynamic scheduling in keeping the solution close to the FBA feasible space, without heavily impacting data-driven performance.

Figure 3 provides a visual comparison of the performance of these methods, focusing on the trade-offs between mechanistic fit and data-driven task performance. The *MINN-scheduler* model demonstrates a balanced performance across both objectives, with a moderate decline in data-driven accuracy but a substantial reduction in mechanistic loss, positioning it much closer to the FBA feasible solution. On the other hand, the models incorporating a mechanistic bound (*MINN-bound* and *MINN-scheduler-bound*) show improvements in mechanistic fit but at a comparable cost in terms of data-driven performance.

**Figure 3:**
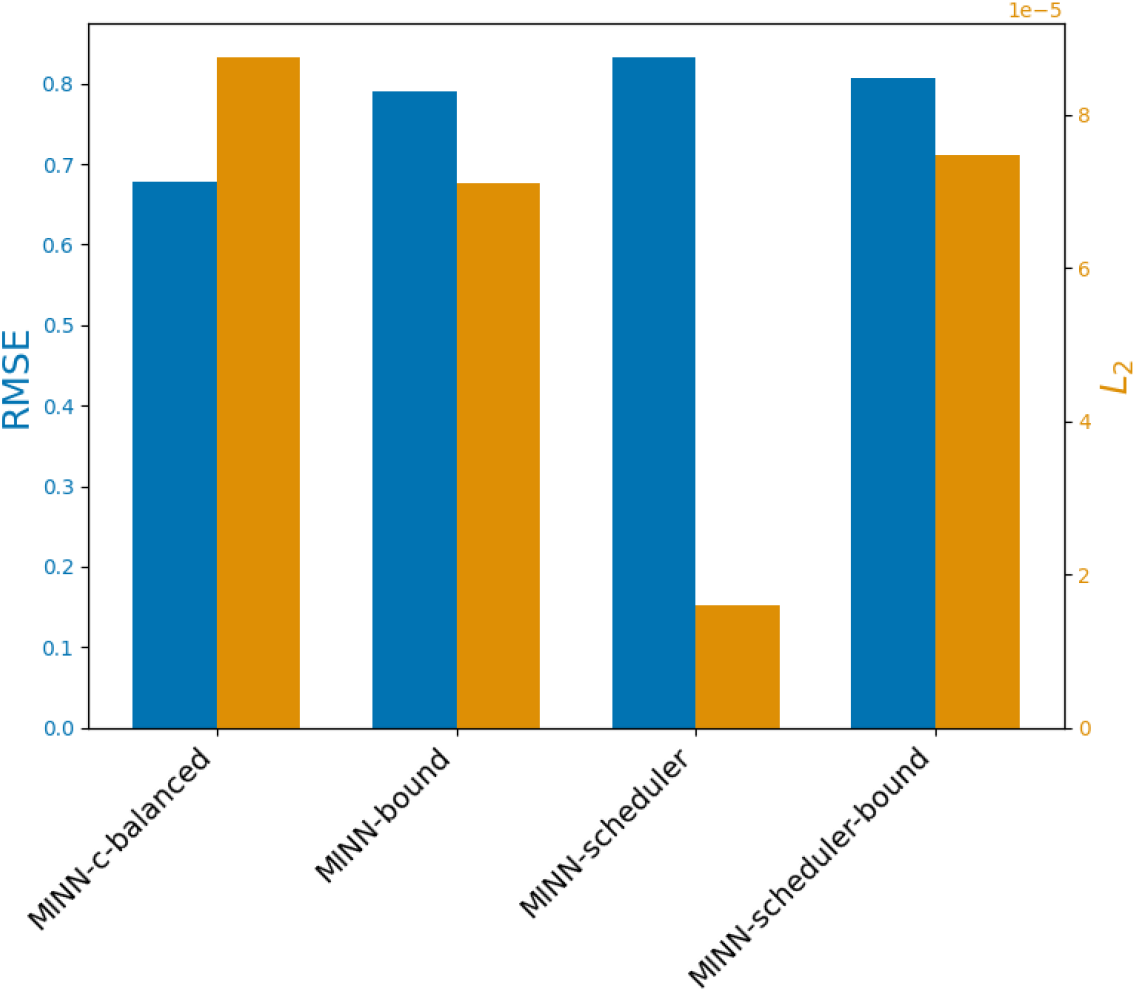
Comparison of different methods based on data-driven task performance (RMSE) and mechanistic fit (*L*_2_ loss), highlighting the trade-off between the two objectives.

These results highlight a clear trade-off between optimizing for mechanistic fidelity and predictive accuracy. While the *MINN-c-balanced* method achieves the lowest RMSE, indicating better performance on the data-driven task, its high mechanistic loss shows that the model prioritizes predictive accuracy over adherence to mechanistic constraints. The models imposing a bound on the mechanistic loss, *MINN-bound* and *MINN-scheduler-bound*, show a even more evident trade-off between these objectives. While they slightly improve the mechanistic fit, this comes at the cost of data-driven performance. In contrast, the *MINN-scheduler* method effectively reduces the mechanistic loss, with only a marginal increase in RMSE.

Overall, these results emphasize the need to carefully select hybrid optimization methods based on the specific priorities of the task. In cases where mechanistic accuracy is crucial, dynamic scheduling methods like *MINN-scheduler* provide a balanced solution, allowing the model to gradually adjust the emphasis between mechanistic fidelity and data-driven optimization.

Additionally, as shown in the third section of Table 3 we tested a configuration of the MINN, called MINN-divided loss, where the the loss is purely data driven (*L*_*MINN*_ = *L*_1_), to check if we could further improve the prediction performance at the cost of *L*_2_. The results show that the improvement in terms of prediction performance is marginal w.r.t. the substantial (four order of magnitude) increase in the *L*_2_ value. This confirms that the regularization effect happens exclusively in the Mechanistic Layer while the *L*_*MINN*_ it is needed to obtain a qualitative flux distribution as output of the MINN.

### 4.3 MINN-reservoir to improve pFBA predictions

In this third group, we present results for the MINN-reservoir method, which extends the use of MINN beyond applications focused solely on prediction. As introduced by Faure et al. [2023], the MINN-reservoir can be used to generate constraints for a mechanistic model in a data-driven manner. In our case, we trained the MINN-reservoir to predict three additional exchange fluxes *(R_EX_co2_e, R_EX_etoh_e and R_EX_ac_e)* which were then used as additional inputs for pFBA. FBA relies on optimization and is generally more accurate in predicting metabolic shifts caused by gene knockouts only after the microbial population has undergone an adaptation period. It is therefore of interest to explore whether the data-driven insights provided by the MINN-reservoir can help overcome this limitation in the case of the newly generated knockout strains from the ISHII dataset. Our baseline consists of a standard pFBA model that uses only the uptake rates of glucose (*R_EX_glc D_e*) and oxygen (*R_EX_o2_e*) as inputs. We compare it with pFBA when it is provided with the same inputs plus the three extra constraints generated by the MINN-reservoir. This setup reflects a realistic use case, in which multi-omics data are available for all samples, while fluxomics data are only partially available. In this context, the MINN-reservoir allows us to estimate missing input fluxes from omics measurements, avoiding the need to experimentally quantify them for every sample.

The advantage of this neural network-based approach is that, once trained, it can generate additional inputs for pFBA based on initial conditions alone, enabling a more informative and automated modeling pipeline. As shown in Table 4, the enhanced version (MINN-reservoir + pFBA) slightly improves the performance in terms of average metrics across the 47 fluxes. However, the most evident benefit is the reduction in the standard deviation, which indicates that the model produces more stable and consistent predictions across the 29 leave-one-out splits.

## 5 CONCLUSION

Faure et al. [2023] introduced a new hybrid architecture, which incorporates a Genome-Scale Metabolic Model in a neural network structure, and used it to predict *E. coli* growth rates in different growth media. In our work, we adapted this framework into a Metabolic-Informed Neural Network, which also uses multi-omics data as input, and tested it with a more challenging task: predicting metabolic fluxes for different E. coli single-gene KO strains grown in minimal glucose medium. The MINN showed improved performance compared to traditional machine learning, and the mechanistic component exerted a regularizing effect on the predictions. We then explored the effect of different components of the architecture on the predictions and their accuracy. Finally, we tested the ability of the MINN to reconcile data and models in a flexible way, even in scenarios where data-driven and mechanistic optimization show a trade-off. To achieve this, we suggested different hybrid optimization strategies.

We chose a naive multi-omics integration approach, such as early concatenation. While the predicting performances are encouraging, as discussed in [Briscik et al., 2024], mixed integration strategies are often to be preferred and they could be tested to further improve the prediction power of a MINN-based method.

Additionally, more work is needed for assessing the interpretability of the flux distribution. As a first step in this direction, in our simulation we kept track of *L*_2_, as a proxy for how much the predicted metabolic profile complies the theoretical assumptions of FBA. Finally, this novel framework has been tested only for *E. coli*, the classical “work-horse” of microbial physiology. We hope the promising result of these works will prompt the creation of suitable datasets to apply this techniques to other microorganisms and to more diverse scenarios, in which secondary metabolism plays a bigger role and the information provided by the GEM is potentially even more effective in complementing the data-driven learning.

Although this was not the most favorable scenario for FBA, the results of the MINN-reservoir highlight its potential as a promising strategy to enable the full integration of FBA into a machine learning framework, effectively combining the advantages of both mechanistic and data-driven approaches.

Finally, with this work we provide a practical guide for choosing the most suitable MINN configuration based on the modeling objective. If the goal is only to achieve high predictive performance, the standard MINN configuration is sufficient. When the aim is to improve the quality of the predicted flux distribution, the optimization strategies presented here can reduce the mechanistic constraints violation. Lastly, if the objective is to enrich mechanistic models with additional inputs, the MINN-reservoir offers a viable solution to generate constraints in a data-driven way, while keeping the structure and interpretability of classical FBA.

## 6 CODE AVAILABILITY

The code used for the analyses presented in this work is available at https://github.com/gabrieletaz/MINN

## 7 ACKNOWLEDGEMENTS

This work was supported by the E-MUSE project funded by the European Union’s Horizon 2020 Research and Innovation Programme [Marie Skłodowska-Curie Grant Agreement number 956126].

