## Supplemental Text for "MINN: A METABOLIC-INFORMED NEURAL NETWORK FOR INTEGRATING OMICS DATA INTO GENOME-SCALE METABOLIC MODELING"

May 14, 2025

### 8 Supplementary Materials

#### 8.1 Hybrid Optimization Strategies for Data-Driven and Mechanistic Integration

Figure 4 shows the bound on mechanistic loss which penalizes solutions that stray too far from the mechanistic loss threshold, encouraging the model to respect mechanistic constraints during training.

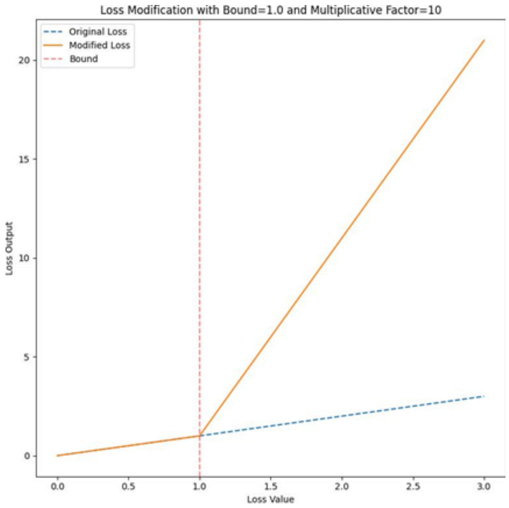


Figure 4: Illustration of the mechanistic loss bound application. The original loss (blue dashed line) remains linear, while the modified loss (orange line) increases steeply after surpassing the bound (red vertical line). This demonstrates how the bound prevents the mechanistic loss from exceeding a set threshold by applying a multiplicative factor beyond this limit.

Figure 5 illustrates how the scheduler dynamically adjusts the weight of the losses over the course of training, allowing the model to optimize both objectives.

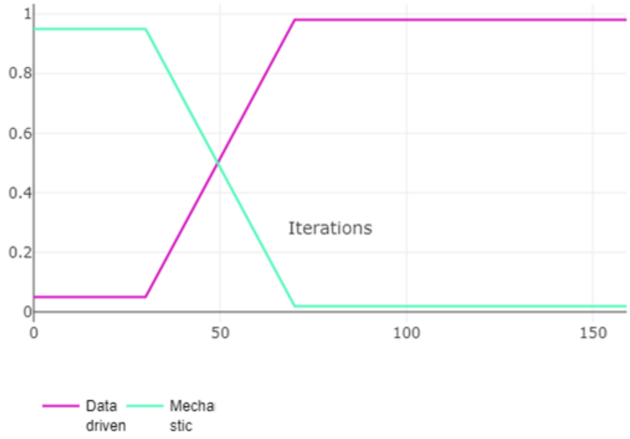


Figure 5: Visualization of the dynamic loss scheduler. The scheduler adjusts the weight of the mechanistic and data-driven losses throughout training, starting with the mechanistic objective and gradually transitioning to prioritize the data-driven objective. This ensures the model initially aligns with mechanistic constraints before focusing on data-driven optimization.

#### 8.2 Other GEMs comparison

Here we present the results of the analysis to explore the role of the GEM in the models' performance. As shown in Table 5, we divided the results in two parts. The first one contains different GEMs in terms of dimension. Here the

GEM is always *E.coli*, but reduced in different ways described in details in the Section 3.2. We also built a MINN with the *E. coli* core model (e_coli_core) from Orth et al. [2010b], to test the smallest version available on BiGG King et al. [2016]. The GEM's dimension can affect the complexity of the NN block in the MINN. Having a layer with many neurons can cause both overfitting and a higher computational time. At the same time, excessively reducing the GEM can decrease the flexibility in the optimization of FBA constraints and make the GEM less representative of the experimental context considered. The first section of Table 5 shows how the FVA reduction have better performances, in terms of metrics and $L_2$, and lower computational time (24h vs 44h) than the full GEM. FBA reduction, instead, performs slightly worse than FVA, but it halves the computational time. On the other hand, the *E. coli* core GEM drastically worse than all the others, making this GEM unfit for this task.

The results show a trade-off between GEM size and computational efficiency. A stricter reduction, such as FBA reduction, shorten the computational time, but at the cost of slightly worse metrics and less reliable flux distribution. While this compromise may not be ideal for small models, it can be helpful for large GEMs, such as yeast or microbial communities, where computational feasibility is critical. In such cases, sacrificing some predictive power in exchange for reasonable runtimes can be a proper trade-off.
Additionally, the poor performance of the E. coli core model highlights the need for an adequate dimension of the GEM, reinforcing the idea that excessively small models may not represent the experimental context of interest.

| | ISHII | | | | |
|---|---|---|---|---|---|
| GEM | $R^2$ | MAE | RMSE | NE | $L_2$ |
| *E.coli* (original) | $0.818 \pm 0.670$ | $0.602 \pm 0.653$ | $1.084 \pm 0.813$ | $0.417 \pm 0.268$ | $4.05 \cdot 10^{-5} \pm 1.57 \cdot 10^{-4}$ |
| *E.coli* (FVA reduced) | $0.950 \pm 0.055$ | $0.473 \pm 0.480$ | $0.678 \pm 0.653$ | $0.272 \pm 0.280$ | $8.75 \cdot 10^{-5} \pm 2.95 \cdot 10^{-4}$ |
| *E.coli* (FBA reduced) | $0.950 \pm 0.048$ | $0.509 \pm 0.518$ | $0.730 \pm 0.719$ | $0.289 \pm 0.295$ | $1.26 \cdot 10^{-4} \pm 2.84 \cdot 10^{-4}$ |
| *E.coli* core | $0.061 \pm 0.099$ | $4.647 \pm 9.959$ | $19.46 \pm 65.30$ | $7.584 \pm 26.70$ | $6.04 \cdot 10^{5} \pm 2.55 \cdot 10^{6}$ |
| *E.coli* (FVA reduced) | $0.956 \pm 0.056$ | $0.512 \pm 0.596$ | $0.759 \pm 0.815$ | $0.285 \pm 0.337$ | $4.27 \cdot 10^{-5} \pm 1.3 \cdot 10^{-4}$ |
| *L.cremoris* (FVA reduced) | $0.954 \pm 0.057$ | $0.546 \pm 0.612$ | $0.801 \pm 0.855$ | $0.304 \pm 0.358$ | $5.24 \cdot 10^{-5} \pm 1.02 \cdot 10^{-4}$ |

Table 5: Performance comparison between different GEMs. Metrics average and standard deviation over 29 leave-one-out splits.

The second part includes a comparison between two GEMs representing two different bacteria, namely *E.coli* and *Lactococcus lactis subsp. cremoris*. The aim of this analysis is to explore how relevant is the nature of the GEM in the MINN architecture. We want to investigate if the regularization that improve the predictive power of the MINN w.r.t. a classical ML approaches is based on a relevant biological information injected in the model through the mechanistic layer, or it's simply a random type of regularization such as Dropout Srivastava et al. [2014]. Since the *L.cremoris* GEM does not have some of the reactions present in the ISHII dataset, in order to have a fair comparison with the *E.coli* GEM, we reduced the number of fluxes to only the one in common between *L.cremoris* and the ISHII dataset. As expected, using a GEM which belongs to another bacteria worsen the prediction performance and also the quality of the predicted flux distribution.

However, the difference in performance between the *E.coli* and *L.cremoris* GEMs is not particularly large. One possible explanation is that the measured fluxes in ISHII dataset belong to the central carbon metabolism, which is highly conserved between both bacteria, reducing the impact of GEM differences. Another factor could be the neural network data-driven component, which may help to compensate for discrepancies between GEMs, reducing their effect on predictive performance. While further investigation is needed, these results suggest that the nature of the GEM plays an important role in the MINN framework. The mechanistic layer likely contributes with biologically relevant information beyond acting as a generic regularization mechanism.
